# Evolutionary rewiring of the dynamic network underpinning allosteric epistasis in NS1 of influenza A virus

**DOI:** 10.1101/2024.05.24.595776

**Authors:** James Gonzales, Iktae Kim, Wonmuk Hwang, Jae-Hyun Cho

## Abstract

Viral proteins frequently mutate to evade or antagonize host innate immune responses, yet the impact of these mutations on the molecular energy landscape remains unclear. Epistasis, the intramolecular communications between mutations, often renders the combined mutational effects unpredictable. Nonstructural protein 1 (NS1) is a major virulence factor of the influenza A virus (IAV) that activates host PI3K by binding to its p85β subunit. Here, we present the deep analysis for the impact of evolutionary mutations in NS1 that emerged between the 1918 pandemic IAV strain and its descendant PR8 strain. Our analysis reveal how the mutations rewired inter-residue communications which underlies long-range allosteric and epistatic networks in NS1. Our findings show that PR8 NS1 binds to p85β with approximately 10-fold greater affinity than 1918 NS1 due to allosteric mutational effects. Notably, these mutations also exhibited long-range epistatic effects. NMR chemical shift perturbation and methyl-axis order parameter analyses revealed that the mutations induced long-range structural and dynamic changes in PR8 NS1, enhancing its affinity to p85β. Complementary MD simulations and graph-based network analysis uncover how these mutations rewire dynamic residue interaction networks, which underlies the long-range epistasis and allosteric effects on p85β-binding affinity. Significantly, we find that conformational dynamics of residues with high betweenness centrality play a crucial role in communications between network communities and are highly conserved across influenza A virus evolution. These findings advance our mechanistic understanding of the allosteric and epistatic communications between distant residues and provides insight into their role in the molecular evolution of NS1.

## Introduction

Unraveling how mutations affect protein function—referred to as sequence-function space—is crucial for understanding the trajectories of protein evolution (1). However, even with advancements in high-throughput techniques like deep mutational scanning, it is nearly impossible to test all possible combinations of mutations in a protein (2). A major challenge is epistasis, which refers to nonadditive interactions between mutations (3, 4). Namely, epistasis makes the effect of a mutation depend on the presence or absence of others. As a result, epistasis exponentially increases the number of potential inter-residue interactions as the number of residues in a protein increases (2).

Studies have shown that epistasis is prevalent and plays a multifaceted role in protein evolution and engineering (3, 5-7). For example, it is a critical factor in predicting immune-evading mutations in viruses, such as SARS-CoV-2 (8, 9). Epistasis is also important in protein engineering, such as enhancing protein thermostability or altering the substrate specificity of enzymes (10-12).

Phenomenologically, epistasis is closely related to protein allostery (13-15). Allostery occurs in response to the binding of a ligand or protein to a site distant from the functional site. Similarly, epistasis involves the communications between mutations located at a distant site (16, 17). Therefore, epistasis and allostery share the similar mechanistic bases (14, 18, 19). Despite significant progress (20-22), however, identifying the mechanism by which ligand binding or mutational effects are propagated to a distant sites across a protein structure remains challenging. This difficulty arises partly due to the intricacies in quantitatively assessing subtle changes in conformation or dynamics at an atomic resolution (23-28).

Here, we examine the molecular basis of epistasis in nonstructural protein 1 (NS1), a major virulence factor of an influenza A virus (IAV)(29). NS1 interacts with various host factors to suppress innate immune responses of host cells. For example, NS1 binds to retinoic acid-inducible gene I (RIG-I) (30), tripartite motif containing 25 (TRIM25) (31), and phosphatidylinositide 3-kinase (PI3K) (32, 33). Notably, the NS1-PI3K interaction is mediated by effector domain of NS1 (NS1ED) and the p85β subunit of PI3K (**Fig 1A**). This interaction is highly conserved across all IAV strains, underscoring its functional importance (32, 33).

**Figure 1.**
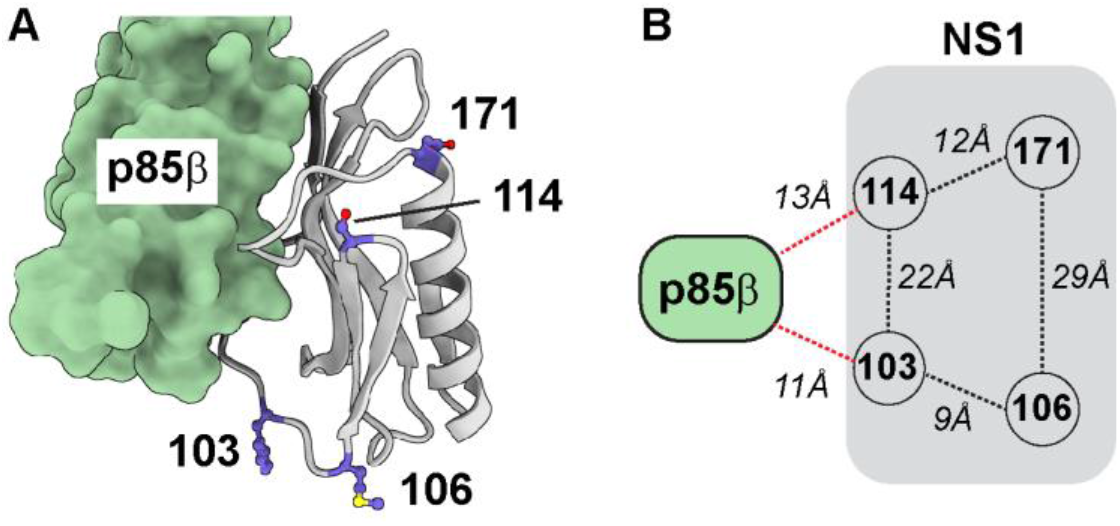
Structure of the NS1-p85β complex (PDB: 6U28). (**A**) The four mutation sites in NS1 (gray) are shown with a ball and stick model. (**B**) Distances between NS1 (shown in gray) and p85β. Dashed lines represent the closest Cα-Cα distances: between mutation sites (shown in black) and between mutation sites and p85β (shown in red).

The NS1ED of the 1918 IAV, which caused the Spanish flu in 1918, binds to PI3K with a high affinity by forming a ternary complex with human CRK (**Fig 1A**) (34-36). Although this ternary interaction increases the virulence of 1918 IAV, NS1s of its descendant strains lost the CRK-binding function (34, 36).

For example, the NS1ED of PR8 strain, which emerged in 1934, does not bind to CRK (34, 37), but harbored four substitutive mutations (**Fig 1A**) that enhance its intrinsic binding affinity to p85β, independent of CRK-binding (21). Therefore, these mutations represent the early trajectory of the adaptive evolution of 1918 NS1 in humans (38). Intriguingly, none of these four mutations are located on the p85β-binding surface, indicating that these mutations increase the binding affinity through long-range effects (**Fig 1B**), i.e., an allosteric mechanism.

Our previous study showed intra-molecular epistatic interactions between strain-specific mutations and p85β-binding residues within NS1ED (hereinafter NS1) (21). However, several important questions remain to be addressed. For example, although it was demonstrated that side-chain dynamics were altered during NS1 evolution (21, 33), it remains unclear how the altered dynamics epistatically change the p85β-binding affinity of NS1s.

This study aims to reveal high-resolution mechanism by which mutational impacts are propagated over long distance. We first characterize the epistatic interactions among the strain-specific mutations occurred between 1918 and PR8 NS1s. We demonstrate how these mutations epistatically alter the structure and dynamics of PR8 NS1 using NMR approaches, with a special emphasis on side chain dynamics. By employing molecular dynamics (MD) simulation along with graph theory-based network analysis (39-41), we identify the epistatic network through which mutational effects are propagated across the NS1 structure. Furthermore, we show how the communications between network communities respond to these mutations. Overall, our study provides a comprehensive picture of how mutational effects are transmitted to distant sites in a protein via dynamic communication network.

## Results

### Allosteric epistasis alters the binding affinity between NS1 and p85β

We measured the binding affinity of NS1 to p85β using biolayer interferometry (BLI). Evolutionary mutations in PR8 NS1 increased the binding affinity to p85β by approximately 10-fold more tightly than 1918 NS1 (**Figs 2A and 2B**). To identify the mutations responsible for this affinity difference, we substituted amino acids in 1918 NS1 with those of PR8 NS1 at the mutation sites and measured their effects on the p85-binding (**Fig 2A and Fig S1**). Notably, two single mutations, S114P and D171A, significantly enhanced the affinity compared to wild-type 1918 NS1. Individually, these two mutations fully recapitulate the affinity of PR8, although neither mutations is located on the p85β-binding surface (**Figs 1A and 1B**). For example, the Cα atoms of residues 114 and 171 are positioned over 11 Å away from p85β in the complex.

**Figure 2.**
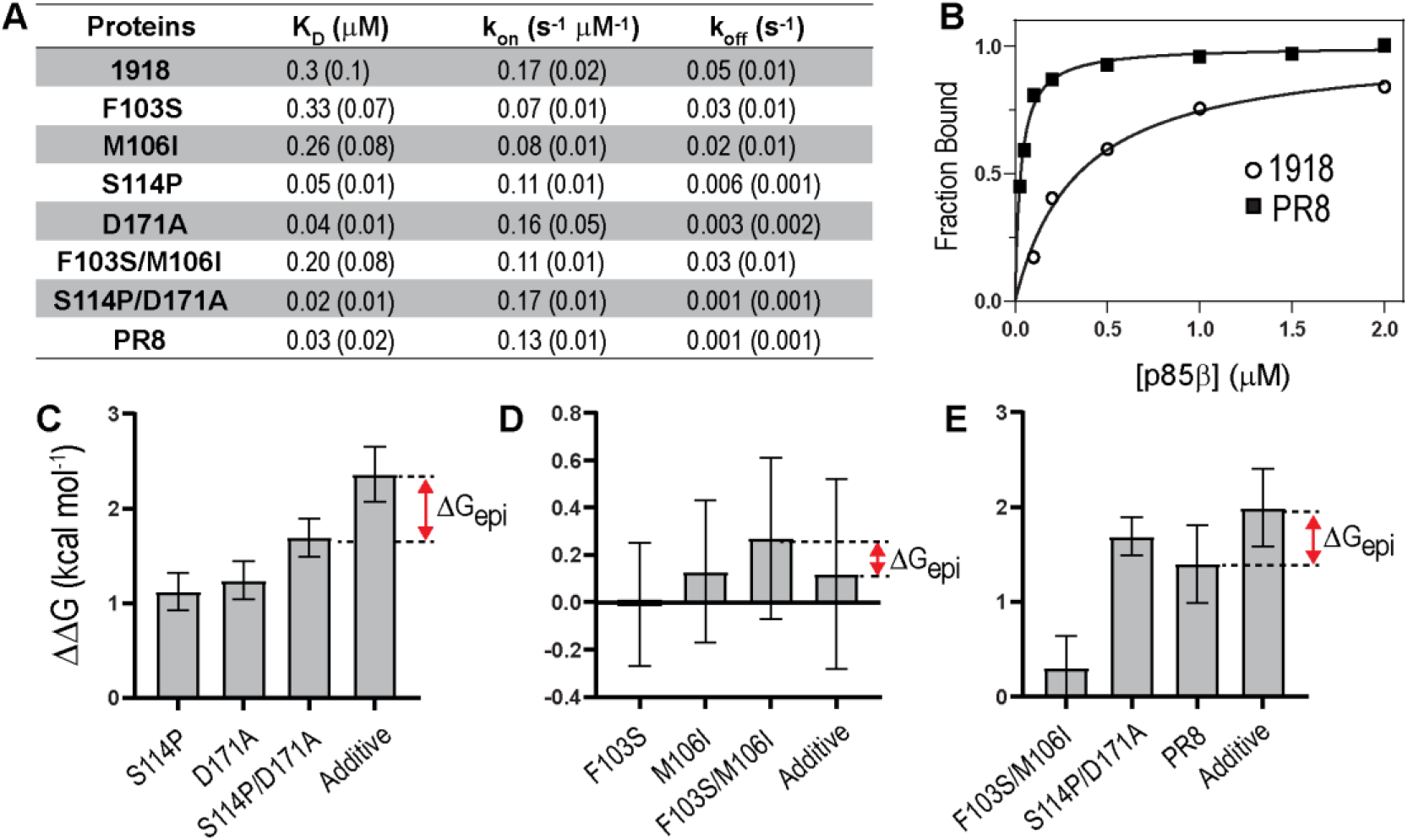
Mutational effects on the NS1-p85β interaction. (**A**) Effects of mutations on binding affinity and kinetics. Numbers in parenthesis are standard deviation of three repeated measurements. (**B**) Representative binding isotherms of wild-type 1918 and PR8 NS1s to p85β. Epistatic interactions (ΔG_epi_) between residues (**C**) 114 and 171, (**D**) 103 and 106, and (**E**) 103/106 and 114/171. PR8 corresponds to quadruple mutations at residues 103, 106, 114, and 171. Additive represents the expected additive effects of mutations.

Furthermore, we observed epistatic interactions between these mutations. Specifically, the effect of double mutation S114P/D171A was 0.7 kcal mol^-1^ less than the expected additive effect of two single mutations (S114P and D171A) (**Fig 2C**). This phenomenon, where the combined effect of two mutations is less positive than the sum of individual effects, is termed the negative magnitude epistasis (1). Although this long-range epistasis could occur by change in global protein stability upon mutations (42), our previous results showed that 1918 and PR8 NS1s have virtually identical thermodynamic stabilities (21), allowing us to exclude the change in protein stability as a contributing factor.

In contrast, two other mutations, F103S and M106I, showed marginal effects on the binding affinity because their effects on k_on_ and k_off_ canceled each other out (**Fig 2A**). This minor mutational effect is not due to their distances to p85β, as they are comparable to those between 114/171 and p85β. Moreover, there was no epistatic interaction observed between F103S and M106I, nor between the double mutations F103S/M106I and S114P/D171A (**Figs 2D and 2E**).

### 1918 and PR8 NS1s have subtle but noticeable structural difference in their free state

Our previous isothermal titration calorimetry (ITC) results indicated that the major driving force of the ΔΔG°_bind_ between the two NS1s is the change in binding entropy. -TΔS°_bind_ was -3.6 ± 0.1 and -5.1 ± 0.4 kcal mol^-1^ for 1918 and PR8 NS1, respectively (21). Namely, PR8 NS1 was associated with a more favorable entropy change upon p85β-binding than 1918 NS1. Given that the two NS1s have virtually identical strucures in the complex with p85β (**Figs S2A and S2B**) (33), the ITC result suggested that PR8 NS1 is accompanied by a less conformational entropy penalty upon binding to p85β (i.e., more favorable ΔG°_bind_). Therefore, we hypothesized that free PR8 NS1 has more bound-like conformation than 1918 NS1.

We sought to test our model which indicates different conformations between the two NS1s in the free state. We initially attempted to compare their crystal structures. However, the crystal structures of the free NS1s showed considerable conformational heterogeneity even for the same NS1 (**Fig S2A and S2C**) (21, 43), preventing meaningful structural comparisons. Thus, we conducted an NMR chemical shift perturbation (CSP) analysis (44, 45). For a comprehensive examination of the mutational effects, we included CSPs of all assigned side-chains ^13^C atoms in addition to backbone atoms because mutations mainly alter side-chain interactions (**Fig S3**). To identify “significant” CSPs, we employed modified z-score (z_M_-score) that is more robust to outliers than the regular z-score (46). The regular z-score is easily affected by the presence of data with large changes, resulting in the exclusion of many residues with intermediate-level changes. Throughout the present study, we defined |z_M_| > 1.5 as a significant difference (**Fig S3**).

The NMR CSP analysis, including side chains, provided new insights into mutational effects that were not identified in the previous study (21). For example, we identified two major clusters of residues showing large CSPs. Cluster-1, centered around mutations at residues 114 and 171 (denoted as 114/171), consists of a substantial number of residues, including solvent-exposed ones. (**Fig 3A**). Cluster-2 was more localized around mutations at residues 103 and 106 (denoted as 103/106) (**Fig 3A**). These two clusters appeared to be separated from each other. This CSP result supports our model wherein the 1918 and PR8 NS1 proteins populate different NS1 structures in their free state.

**Figure 3.**
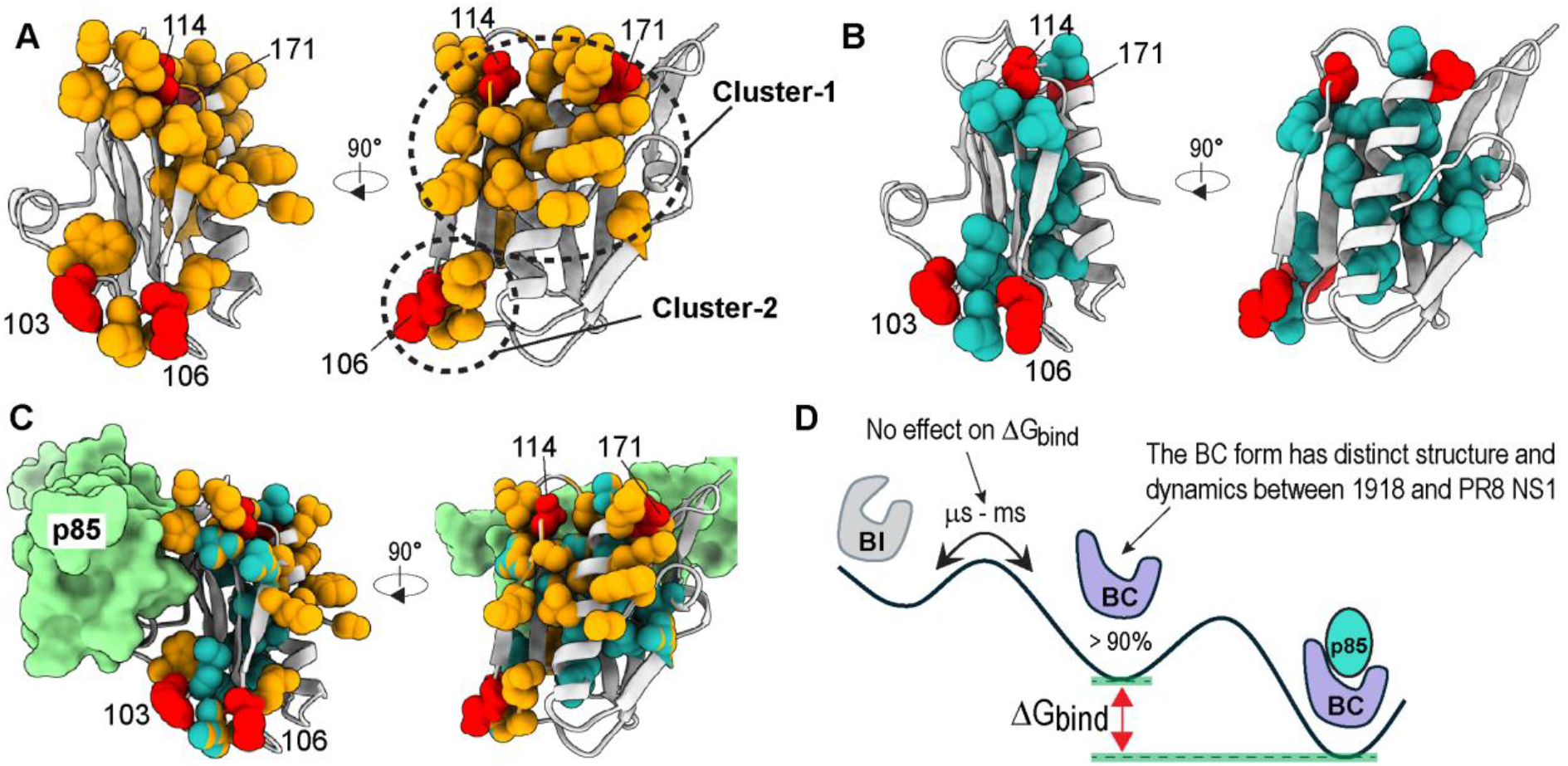
Mutational effects on NS1 structure and dynamics. Residues with a significant NMR (**A**) CSP (z_M_ > 0.5) and (**B**) ΔS^2^_axis_ (> 0.05) are shown as orange and sea green spheres, respectively. (**C**) Residues in (A) and (B) are superimposed. Spheres with a split color represent residues with signficant CSP and ΔS^2^_axis_ . Red spheres represent mutation sites. (**D**) Energy landscape of the NS1-p85β interaction representing that NMR-detected changes in structure and dynamics correspond to the BC form in the free state. The BI-BC dynamics in slow μs – ms timescales have no impact on binding free energy to p85β (ΔG_bind_).

We further examined the role of side-chain dynamics in the epistatic effect on the p85β-binding affinity of NS1s. To do this, we turned to the measurements of the NMR methyl-axis order parameters S^2^_axis_, which quantitatively represent the intrinsic flexibility of individual methyl-moieties of side-chains. We previously employed ^2^H relaxation approaches (47) to measure S^2^_axis_ values of NS1s (21). However, a considerable number of methyl-containing residues were excluded from the analysis because of low signal sensitivity and limited spectral resolution of the ^2^H relaxation experiments. To overcome the limitation, in the present study, we employed highly sensitive intramethyl ^1^H-^1^H dipolar cross-correlated relaxation (CCR) rate approach to measure S^2^_axis_ values of I, L, and V residues (48, 49).

We first assessed the consistency of the S^2^_axis_ values measured by the ^2^H- and CCR-derived approaches using the residues analyzed by both methods. The S^2^_axis_ values showed excellent agreement between the two methods (**Fig S4**). This allowed us to combine the two datasets, incorporating A, M, and T from the previous study (21) with I, L, and V. Furthermore, the root mean squared error (RMSE) between the ^2^H and CCR derived S^2^_axis_ values was 0.04, enabling robust uncertainty estimation for S^2^_axis_ values. Based on the RMSE, we defined a significant difference in S^2^_axis_ values (ΔS^2^_axis_ ) between 1918 and PR8 NS1s as > 0.05.

Next, we examined the spatial distribution of residues showing a significant ΔS^2^_axis_ (> 0.05) between two NS1s (**Fig 3B**). Overall, the dynamically altered residues were located near mutation sites and in the hydrophobic core of the NS1 structure. Notably, many residues in the hydrophobic core showed altered side-chain dynamics, although none of the mutations are located in the hydrophobic core. These residues appear to be networked by continuous physical contacts between side chains. Many of them were not identified by CSP analysis, especially residues in the hydrophobic core (**Fig 3C**), indicating that these residues undergo minor conformational changes.

Moreover, when considering all residues with significant CSP or ΔS^2^_axis_ collectively, it became more evident that mutations of residues 114/171 extensively affected the NS1 structrure and dynamics (**Fig 3C**). Upon visual inspection, we also observed two clusters of affected residues, similar to those identified by CSP analysis, with only marginal connection between the two clusters (**Fig 3C**). This observation is consistent with the epistatic pattern where residues 114/171 show no epistatic interaction with residues 103/106 (**Fig 2E**). These NMR data also explained why S114P and D171A have more impacts on the p85β-binding affinity than F103S and M106I. Cluster-1 included some residues directly interacting with p85β, while none of cluster-2 interact with p85β (**Fig 3C**). Given its higher affinity for p85β, the NMR data suggested that PR8 NS1 adopts a conformation more similar to the p85β-bound state than 1918 NS1.

It is worth mentioning that NS1 in the free state populates two alternative conformations, p85β-binding incompetent (BI) and a binding competent (BC) form (**Fig 3D**) (33). The BC conformer is the major species, corresponding to 90 – 99 % of the entire population in the free state (21). Thus, our NMR data indicated the mutations altered the conformation and dynamics of BC forms. This result is also consistent with the ITC data, where the intrinsic dynamics of the BC form of PR8 NS1 differ from those of 1918 NS1.

### Network analysis reveals long-range communication pathways in 1918 and PR8 NS1s

It is possible that the mutations altered the dynamics of more residues than were probed by NMR, since only methyl-containing residues were included in the NMR S^2^_axis_ analysis. To complement the NMR result and gain further insight into how the mutational effects were propagated across the NS1 structure, we employed atomistic MD simulations in combination with network analysis.

We first assessed the MD simulation results by comparing the calculated backbone NH order parameters (S^2^_NH_) with the previously reported NMR-derived values (21). Overall, the results of two approaches showed reasonable agreement: Pearson correlation coefficients (r) were 0.79 and 0.76 for 1918 and PR8 NS1s, respectively (**Fig S5**).

To analyze the dynamic inter-residue interactions in NS1s, we applied graph theory-based network analysis (41, 50). Briefly, the dynamic residue interaction network (DRIN) analysis represents each residue as a node connected to other nodes by edges representing the correlative dynamics between nodes (50). In our analysis, edge lengths were weighted by generalized dynamic cross-correlation (DCC) coefficients to account for non-linear correlations between residues (**Fig S6**)(51), which are considered critical for allosteric conformational changes (52, 53). Thus, the length of an edge decreases as the two nodes are connected by stronger correlative dynamics.

In the network analysis, the shortest path (SP) and shortest path length (SPL) represent the most efficient communication route and its communication efficiency, respectively, between two nodes in the network (41, 50). The average SPL, also known as the characteristic path length (CPL) (41, 54), indicates the overall connectivity or efficiency of communication between residues. PR8 has a slightly shorter CPL than 1918 NS1s (1.6 for 1918 vs. 1.4 for PR8) (**Fig S7**). Interestingly, however, some mutation sites showed significantly larger changes in SPL between the two NS1s. For example, the SPL between residues 114 and 171 decreased significantly in PR8 NS1 compared to 1918 NS1 (ΔSPL_1918-PR8_ = 0.71), whereas other mutation pairs do not (**Fig 4A**). This observation is consistent with the pattern of epistasis (**Figs 2C-2E**), which suggests that the SPs represent epistatic pathways.

**Figure 4.**
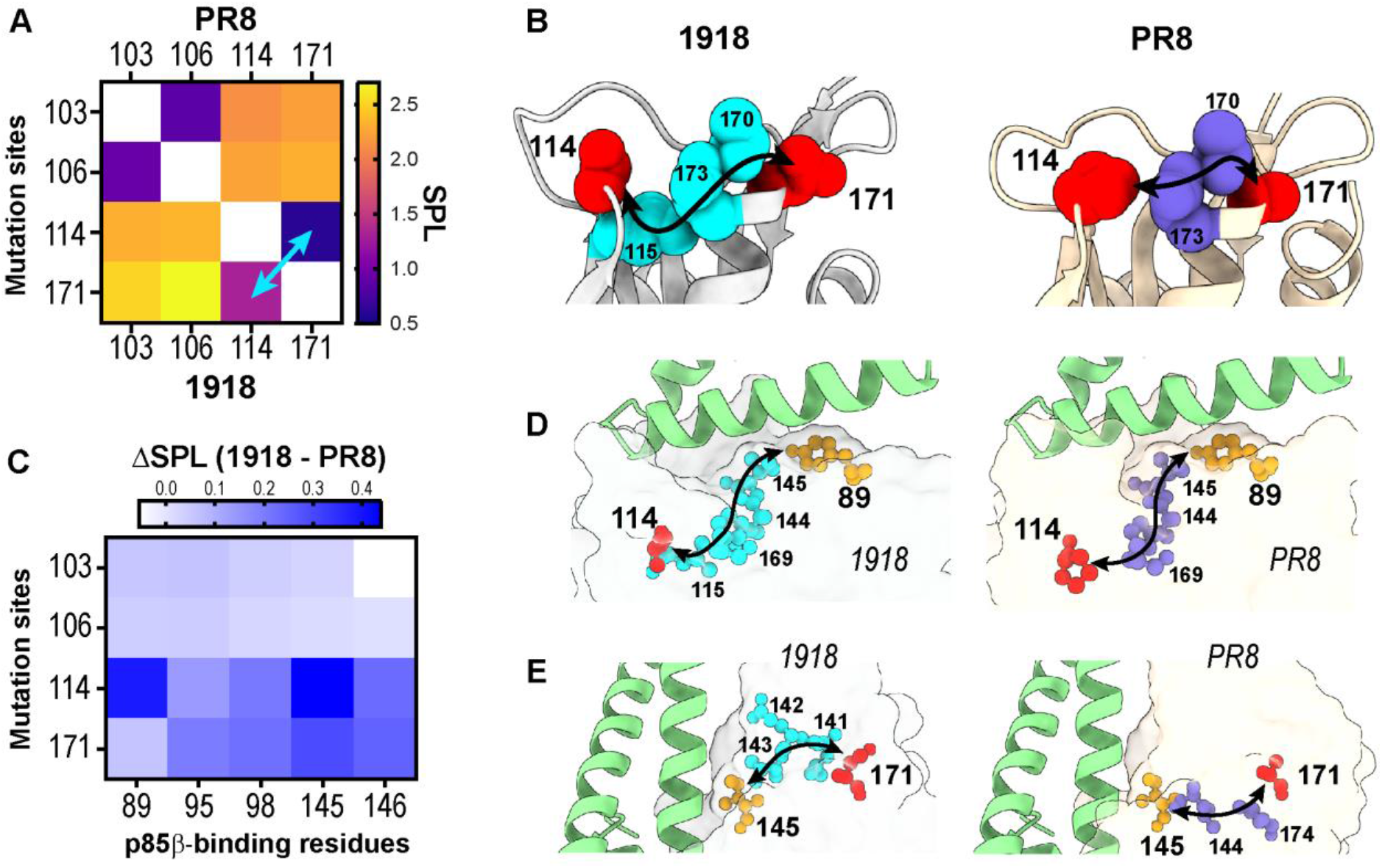
The network analysis of mutational effects. (**A**) Heat map of SPL between mutation sites. The upper and lower triangles of the heat map are SPLs in PR8 and 1918 NS1s, respectively. The cyan arrow indicates the largest SPL change between residues 114 and 171 in two NS1s. (**B**) The shortest pathways (SPs) connecting residues 114 and 171 in 1918 and PR8 NS1s. (**C**) Heat map showing the normalized difference in SPLs between mutations and p85β-binding residues. The (**D**) 89 – 114 and (**E**) 145 – 171 pathways in 1918 and PR8 NS1s. Red and gold spheres represent mutation sites and p85β-binding residues. p85β is shown as a green cartoon. The SPLs in panels A and C are provided in Figure S8.

The change in SPLs could be due to rewired SPs, as observed in allosteric pathways between distant sites (55, 56). Indeed, we noted that the SP between residues 114 and 171 differ in the two NS1s. Specifically, residue 115 was skipped in the 114-171 pathway in PR8 NS1, reducing its SPL compared to that in 1918 NS1 (**Fig 4B**). These changes in SP and SPL may underlie the epistatic effect between the two residues.

To understand how the mutations allosterically affect the p85β-binding affinity, we analyzed SP and SPL of the mutation sites with key p85β-binding residues in 1918 and PR8 NS1s. Two residues, 114 and 171, showed noticeable changes in SPLs, while other mutation sites showed only marginal changes (**Fig 4C**). For instance, residue 114 exhibited a large change in SPL with residue 89, the most critical p85β-binding residue (57). The 114-89 pathway was preserved in both NS1s except for that residue 115 was excluded in the pathway in PR8 NS1, while it is included in 1918 NS1 (**Fig 4D**). In contrast, residue 171 was connected to the p85β-binding residues through distinct pathways in the two NS1s (**Fig 4E**). These results exhibited diverse ways in which mutations rewire the communication between residues.

Notably, the pathways connecting residues 114 and 171 to the key p85β-binding sites include residues that exhibited significant NMR CSP and/or ΔS^2^_axis_ between the two NS1s (**Fig S8**). Importantly, the initial two or three residues in all pathways from residues 114/171 to the core p85β-binding residues were the ones identified by NMR. This result suggests that conformational or dynamics changes near mutation sites are crucial in rewiring the communication networks to the p85β-binding surface. Consequently, mutational effects on proximal residues critically influence the rewiring of long-range communication pathways.

### Mutational effects on network community underpin epistasis in NS1

The network community analysis can reveal the spatial organization of residues according to their correlated dynamics (41, 52) and how a mutation can influence the structure and dynamics of other residues within the same community (55). Hence, the rearrangement of network communities can represent allosteric communication between communities (58, 59), offering complementary perspectives to approaches that focus on individual SPs.

Overall, both NS1s displayed similar, but not identical, geometric distributions of network communities across their structures (**Figs 5A, 5B, and Table S1**). The p85β-binding surface consists of three communities to which residues 114/171 belong (**Fig 5C**). Therefore, mutations at these two residues could alter the dynamics of other residues in the same community that forms the p85β-binding surface, thereby changing the binding affinity. Indeed, NMR data showed that mutations alter the conformation and dynamics of many residues within the communities to which they belong (**Fig 5D**). These results also provide insight into the epistatic communication between residues 114 and 171. Namely, their mutations affect the same community in 1918 NS1, potentially leading to crosstalk between the mutational effects and resulting in epistatic interactions.

**Figure 5.**
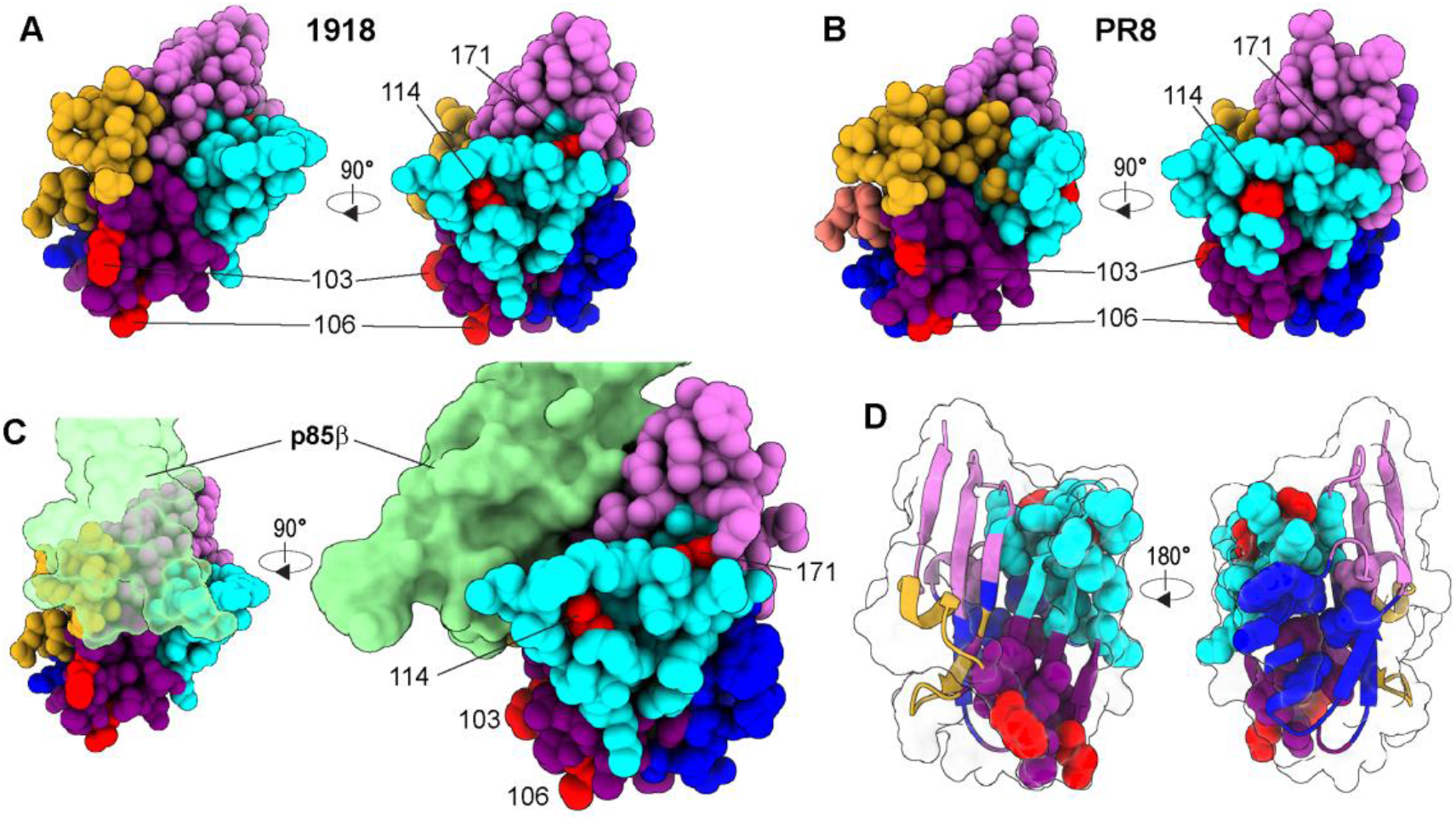
Evolutionary change of network community. Network communities in (**A**) 1918 and (**B**) PR8 NS1s. Spheres in different communities are shown in distinct colors. (**C**) The same NS1 structure as shown in (A) with surface representation of bound p85β (green). Only three communities (shown in gold, orchid, and cyan) interact with p85β. Residues 114 and 171 belong to cyan community. (**D**) The residues with a significant NMR CSP or ΔS^2^_axis_ between 1918 and PR8 NS1s are depicted as a sphere model. The color of these residues correspond to the community to which they belong. The color schemes for communities are the same as in (C). The majority of these residues belong to the same community with mutation sites (red spheres).

Residues 103 and 106 belong to a community that is not a part of the p85β-binding surface (**Fig 5C**). The geometric position of this community might explain why mutations at residues 103 and 106 have only marginal effects on the binding affinity. However, this does not mean these mutations have no impact on other residues in the community. Indeed, the F103S and M106I mutations also perturbed the conformation and dynamics of multiple residues in the community to which they belong, as revealed by NMR data (**Fig 5D**). These perturbations were limited within the community that does not contribute to the p85β-binding surface. Thus, NMR data provide critical experimental validation for the coupled dynamics that underpin the network community analysis.

The network analysis is also useful to identify critical residues that control the information flow within the dynamic network. To identify these critical residues, we computed the betweenness centrality (C_B_) of individual residues of NS1. C_B_ is the measure of the frequency with which a node lies on the shortest paths between other nodes (60). The residues with high C_B_ are often reported to play critical roles in protein stability and allosteric communications (56, 61).

We noted that the eight of the top 10 C_B_ residues were preserved between 1918 and PR8 NS1s (**Fig 6A**), indicating their evolutionary importance. Moreover, nine of the top 10 C_B_ nodes in NS1 are hydrophobic residues. Presumably, their high packing density with multiple residues results in correlated motions among them. The inverse correlation between C_B_ and surface accessibility was also observed in other proteins (60). Mutation of high C_B_ residues is proposed to significantly alter the allosteric network (60, 62). Therefore, the hydrophobic residues with high C_B_ may be important for maintaining the global structural/dynamic network of NS1.

**Figure 6.**
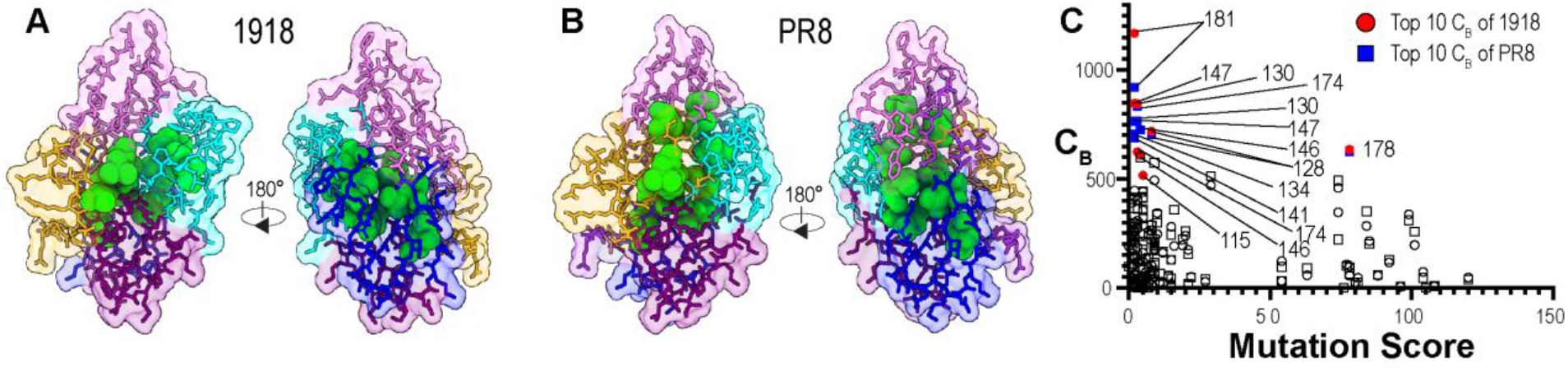
Inter-community communications. (**A**) 1918 and (**B**) PR8 NS1s with top 10 betweenness centrality (C_B_) residues (shown as green spheres). The green residues are located in the community interfaces. Other residues are depicted using a stick model. The residues in different communities are represented by different colors. (**C**) Relationship between C_B_ and mutation frequency during NS1 evolution. The mutation score was calculated using NS1 sequences of human H1N1 IAVs (n = 2304). Circles and squares correspond to residues of 1918 and PR8 NS1s, respectively. Top 10 C_B_ residue numbers are shown.

We also noted the high C_B_ residues of NS1 are located at the interfaces between network communities, indicating their critical role in inter-community communications (**Figs 6A and 6B**) (63). Significantly, our NMR study showed that five of the top 10 high C_B_ residues underwent significant changes in side-chain dynamics between 1918 and PR8 NS1s. Considering their geometric feature, we conjecture that the altered dynamics is associated with adaptation of the network upon mutation, which enables other residues to explore new communication routes. To our knowledge, this is the first experimental observation where high C_B_ residues alter their side chain dynamics in response to mutations.

## Discussions

The mechanism by which mutations reshape the conformational energy landscape is critical for understanding protein evolution trajectories and has significant impacts on protein design and engineering. Despite recent advances in AI-based prediction of protein structures, our knowledge of how mutational effects propagate across a protein structure remains incomplete (64, 65). This limitation hinders our mechanistic understanding of how epistasis reshapes the energy landscape.

Strain-specific mutations harbored in PR8 NS1 increased its binding affinity to p85β relative to its ancestor, 1918 NS1, although none of the mutations are located on the binding surface. Here, our deep mechanistic study revealed how the mutations rewired the dynamic network underlying their epistatic interactions and allosteric effects on the p85β-binding affinity. Notably, we focused on how side chain dynamics contribute to the networked relay of inter-residue communications across the protein structure. Although mutations mainly alter side chain interactions, surprisingly little is known about how side chain networks propagate mutational effects.

Intramolecular epistasis shares a similar mechanistic basis with protein allostery because both phenomena depend on the long-range communication within a protein (14, 19). A combination of NMR and computational approaches has revealed new perspectives on allosteric mechanisms (52, 58, 66). For example, the violin model was proposed based on the findings that a community of residues can share the coupled dynamics. The redistribution of these coupled dynamics upon binding of an effector underlies the propagation of conformational dynamics (59). This model contrasts with the traditional domino-like model, which relies on specific pathways connected by physical contacts of residues (58, 67). Hence, we examined whether these models are also applicable to understanding the mechanisms of epistatic effects.

In the present study, we observed that the propagation of mutational effects exhibits traits of both models. For example, significant mutational effects of residues 114/171 on the p85β-binding affinity can be explained by the network community model. Both residues in 1918 NS1 belong to the same network community, which constitutes the major p85β-binding surface. Thus, mutation of either residue could affect the other through coupled dynamics in the community, resulting in epistatic communication between the two mutations. Our observation of the community rewiring around the p85β-binding surface is also suggestive of the violin model(59).

At the same time, shortest paths (SP) between mutations were identified, and these pathways were connected by contiguous contacts of side-chains, as implicated in the domino model. Moreover, our NMR results indicated that the SP based on physical contacts is highly plausible due to densely populated side-chains. The patterns of changes in SP and SPL between 1918 and PR8 NS1s are consistent with the experimentally observed epistatic interactions, supporting the domino model. Long-range propagation of altered side-chain dynamics through contiguous residues was also observed in other proteins(68-70). Therefore, the violin and domino models provide mechanistic insights from different angles into how mutational effects are propagated through network of inter-residue dynamics.

The betweenness centrality (C_B_) is used to identify residues acting as bridges by controlling the information flow in the network (71). These bridge residues in NS1s are hydrophobic residues located at the interface between network communities (**Fig 6A**). Our NMR data showed that their side-chain dynamics significantly differ between 1918 and PR8 NS1s, and their altered dynamics are intimately related to rewiring the communication (i.e., epistatic) pathways in response to mutations.

These findings raised a question about the evolution of these bridge residues. If they play critical roles in controlling the communication network and epistasis, are they evolutionary conserved? Our study showed that high C_B_ residues are well-conserved between the two NS1s. However, PR8 is considered an early descendant of 1918 IAV, separated by a short evolutionary time period (72). So, we calculated the amino acid conservation of NS1s of all human H1N1 IAVs emerged between 1918 and 2020 (n = 2304), then compared the C_B_ scores and sequence conservation scores of individual residues. Interestingly, nine of the top 10 highest C_B_ residues were highly conserved along the NS1 evolution (**Fig 6C**). Although V178 was the only exception with low-level conservation, it was due to isosteric mutations with I178. Therefore, majority of bridge residues are well-conserved across NS1 evolution, supporting their critical role in maintaining structure and dynamics during evolution. However, it should also be noted that not all evolutionary conserved residues are bridge residues (**Fig 6C**).

It is worth mentioning that residue interaction network can be created from various features such as residue contacts in a static structure, mutational effects on functional activity, amino acid sequence conservation, or conformational dynamics in different timescales (73-76). Each of these approaches will reveal distinct network features associated with different functional properties. Therefore, it will be of important to develop the analysis of multi-scale, multi-layer networks to integrate these features. Overall, our study shed light on the comprehensive picture of how mutational effects are transmitted to distant sites via networked communications.

## Materials and Methods

### Protein sample preparation

All proteins were expressed with the N-terminal His_6_ and SUMO tags in BL21 (DE3) E. coli cells and purified as described in detail elsewhere (21). All purified proteins were > 95% pure; the purity of protein samples was assessed by sodium dodecyl sulfate polyacrylamide gel electrophoresis.

### Biolayer interferometry

The binding of surface-immobilized NS1 to p85β was measured at 25 °C using an Octet RED biolayer interferometry (Pall ForteBio). The N-terminal His_6_ and SUMO-tagged NS1 proteins were used for immobilization. The buffer was 20 mM sodium phosphate (pH 7), 150 mM NaCl, 1% bovine serum albumin, and 0.6 M sucrose (77). All reported values are the average and standard deviation of three repeated measurements. Association and dissociation data were fit using single exponential growth and decay functions, respectively, using GraphPad Prism (ver. 9).

### NMR CSP

Assigned chemical shifts were obtained from the Biological Magnetic Resonance Bank under accession code 52467 for 1918 NS1 and 52466 for PR8 NS1 (21). The NMR CSP per residue was calculated using the following equation.

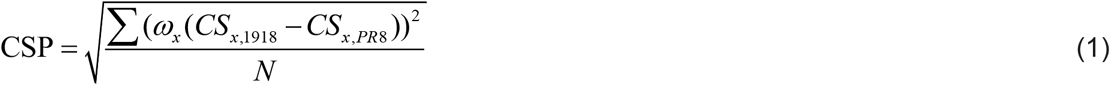

where ω_x_ represents the weighting factor accounting for the differences in the gyromagnetic ratios of different types of nuclei. CS represents a chemical shift. N represents the number of nuclei for individual residues included in the calculation.

### NMR side-chain order parameters

Highly detuerated, ^13^CH_3_-methyl labeled samples were prepared as described elsewhere (78). All NMR samples were prepared in a buffer containing 20mM sodium phosphate (pH 7), 80 mM NaCl, 1mM TCEP, 1mM EDTA, and 99.9% ^2^H_2_O. Intramethyl ^1^H-^1^H dipolar cross-correlated relaxation rates (48) for I, L, and V residues of 1918 and PR8 NS1s were measured by using the following sets of relaxation delays: 4, 8, 16, 20, 24, 28, 32, 37, 41, 44, and 48 ms at 25 °C on Bruker 800 MHz NMR spectrometer. Uncertainties of the relaxation parameters were estimated using duplicated measurements.

Side-chain methyl axis order parameters S^2^_axis_ were calculated using the following equation (48).

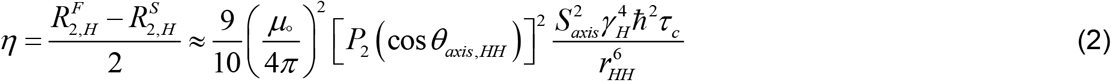

where R^F^_2,H_ and R^S^_2,H_ are the fast and slow relaxation rates, respectively, μ_0_ is the vacuum permittivity constant, θ_axis,HH_ is the angle between the methyl 3-fold axis and a vector connecting a pair of methyl ^1^H nuclei, γ_H_ is the gyromagnetic ratio of a proton spin, r_HH_ is the distance between pairs of methyl protons, and τ_c_ is the global protein tumbling time in ^2^H_2_O.

### Sequence conservation

All human H1N1 IAV sequences (2,304 nonduplicated sequences) were obtained from the NCBI influenza research database (https://www.fludb.org/brc/home.spg?decorator=influenza). The sequences were aligned using MUSCLE (Multiple Sequence Comparison by Log-Expectation) algorithm.

### MD simulations

We prepared two independent systems for the MD simulations for the PR8 (PDB ID: 3O9S (79)) and 1918 (PDB ID: 6U28 (33)) NS1 proteins. Both NS1s include residues from A86 to S205. Structures were prepared and analyzed using CHARMM (80, 81) version c48a1 with the param36 all-atom force field(82), and the TIP3P explicit water model (83). Production dynamics runs were performed using OpenMM version 7.6.0 (84). Each system was independently simulated for a total time of 1 μs.

Both NS1 systems were solvated in a cubic water box with at least 13 Å between the protein and the boundary of the box. Each system was neutralized by adding sodium (Na^+^) and chlorine (Cl^-^) ions, to achieve a net charge of 0 and a concentration of 50 mM. The neutralized systems were then used in a 6-stage energy minimization procedure. During the first stage, 500 steps of the steepest descent (SD) and 1500 steps of the adopted basis Newton-Raphson (ABNR) minimization algorithms were applied on the solvent atoms. The five following stages applied 200 steps of SD and ABNR to the solvent and protein atoms. During these five stages, harmonic constraints were applied to the heavy atoms in the protein backbone and side-chains. The harmonic constraints were initially set to 40 kcal/[mol·Å^2^] and 20 kcal/[mol·Å^2^] for the protein backbone and side-chain atoms, respectively. The constraints were gradually reduced by 10 kcal/[mol·Å^2^] on the backbone atoms and 5 kcal/[mol·Å^2^] on the side-chain atoms until no constraints were applied in the final stage of the minimization. After the energy minimization procedure, both systems were heated from 30 K to 300 K over 100 ps with harmonic constraints of 10 kcal/[mol·Å^2^] applied to the heavy atoms in the protein backbone. The systems were then were equilibrated at 300 K for 400 ps with harmonic constraints of 5 kcal/[mol·Å^2^] applied to the backbone heavy atoms and then with reduced constraints of 0.001 kcal/[mol·Å^2^] applied to the C_α_ atoms for 2 ns. Production dynamics runs were carried out for 1 μs for both NS1 systems. Coordinates were saved every 10 ps, yielding 10,000 frames for every 100 ns or 100,000 frames in total.

### Contact network analysis

The contact network utilized the generalized dynamic cross-correlation (DCC) (55) coefficients to compute the edge lengths between nodes in the graph. The positions of the center of mass of the side-chain heavy atoms were used to calculate the DCC coefficients. To minimize the effects of irrelevant and coincidental correlations, we removed edges in the graph (i.e. correlations between two residues) that did not have physical contacts during the simulation. Physical contacts were identified in our trajectories by using the COORdinates HBONd and COORdinates DISTance RESIdue commands available in CHARMM. Hydrogen bonds are identified with a donor-acceptor distance cutoff of 2.4 Å and nonpolar interactions are classified as two residues within a 3.0 Å cutoff, each with a partial charge of less than 0.3ε (ε = 1.6·10^-19^ C). Only edges between residues with a contact occupancy of greater than 70% were retained, with contact occupancy defined as the percentage of frames where the contact was formed. These “filtered” DCC coefficients (C_DCC_) were then used for determining the lengths of the edges in the graph for our analysis. The NetworkX package (85) in Python was used to perform the graph theory analysis. To find the shortest paths between two nodes in the graph, Dijkstra’s algorithm was used, where the distance between nodes was computed as D = -ln(|C_DCC_|). Since the DCC coefficients are bound between [-1, 1], residues who’s motion is either fully correlated (C_DCC_ = 1) or anti-correlated (C_DCC_ = -1) will have edge lengths of 0 and residues with uncorrelated motion (C_DCC_ = 0) will have large edge lengths. The generalized modularity measures the level of organization within a graph and ranges from [-1/2, 1] (86), graphs with higher generalized modularity are more organized when compared to random noise. Thus, communities were identified using the Clauset-Newman-Moore greedy modularity maximization algorithm (87). This method was chosen as it identifies communities that maximize the generalized modularity of the graph.

## Acknowledgements

This work was funded by NIH grants R01GM127723 (J-H.C.), R35GM152007 (J-H.C.), and the Welch Foundation grant A-2028-20230405 (J-H.C.).

## Notes

### Competing Interest Statement

The authors have declared no competing interest.

