## Supplementary Information for "Evolutionary rewiring of the dynamic network underpinning allosteric epistasis in NS1 of influenza A virus"

Figure S1

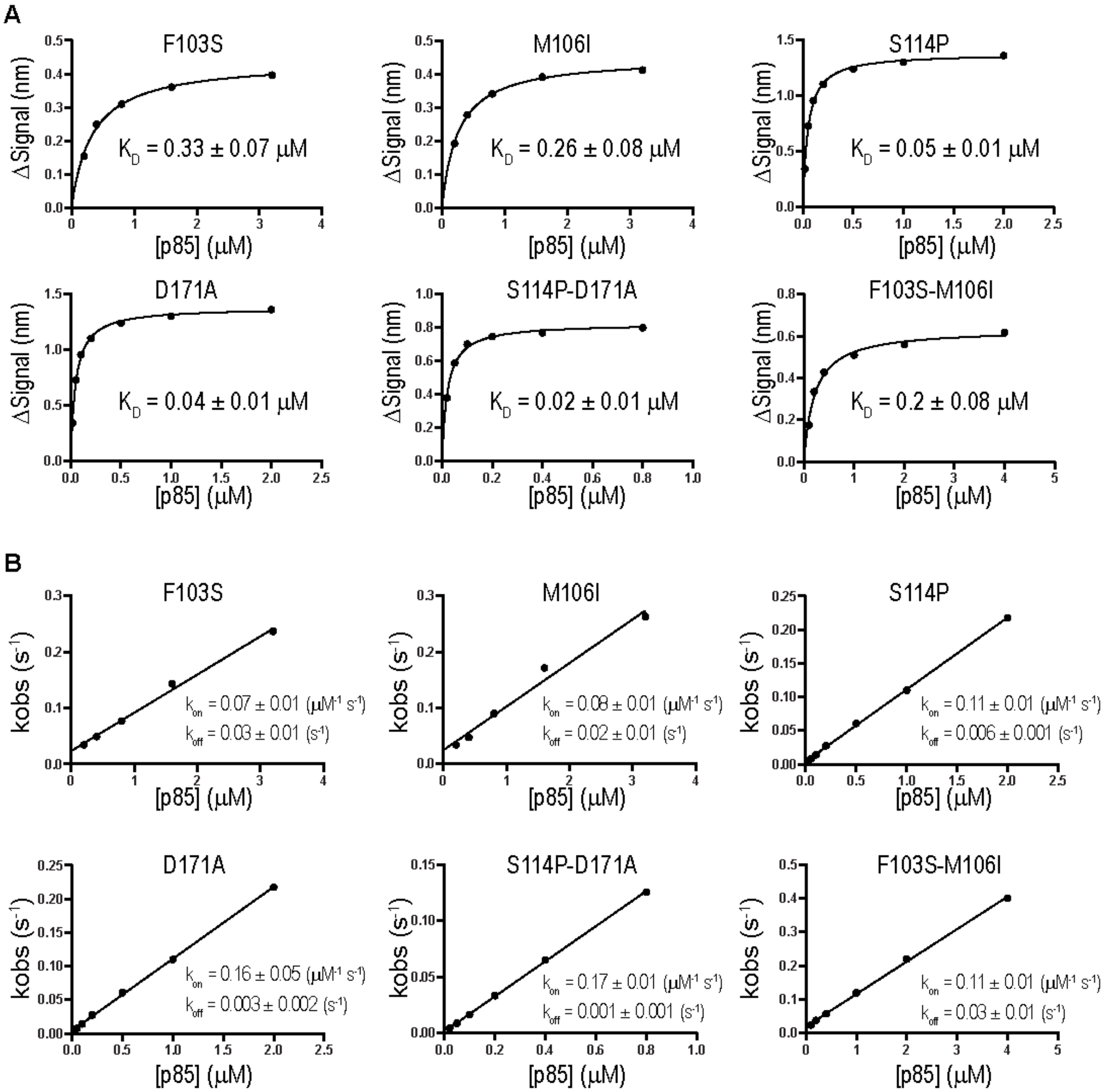


**Figure S1.** Mutational effects of NS1 measured by biolayer interferometry (BLI). (**A**) Representative equilibrium binding isotherms of mutant NS1s and p85β. (**B**) Plots of $k_{obs}$ versus [p85] for mutant NS1s. Numbers after ± symbol represent standard deviations of three repeated measurements.

Figure S2

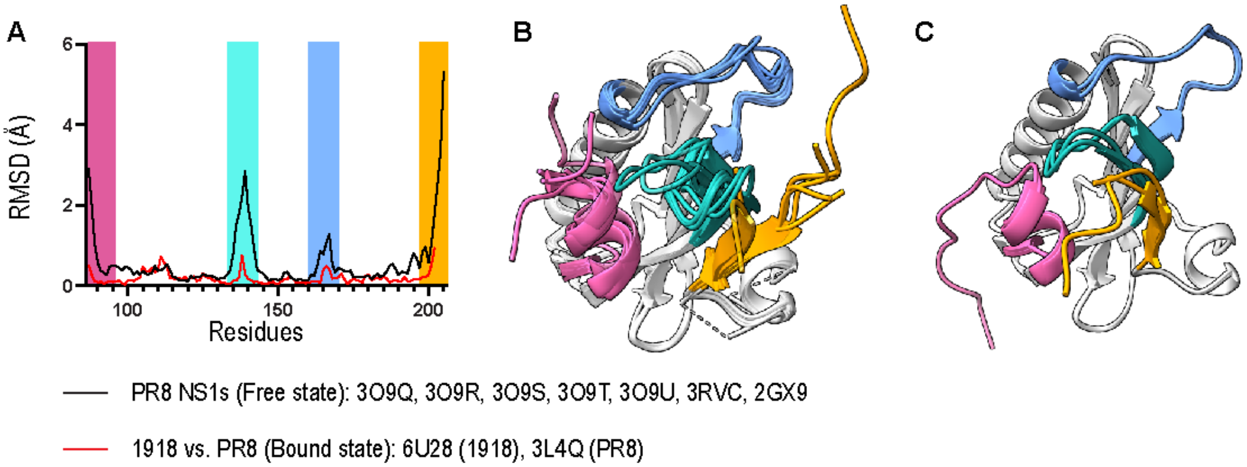


**Figure S2.** Structural heterogeneity of NS1. (**A**) The Cα RMSD of all free PR8 NS1 crystal structures (black line). The Cα RMSD between 1918 and PR8 NS1s bound to p85β (red line). Superimposed crystal structures of (**B**) PR8 and (**C**) 1918 NS1s. The conformationally heterogeneous regions are highlighted in different colors across all panels. The PDB IDs of the superimposed structures are listed at the bottom of panel (A).

Figure S3

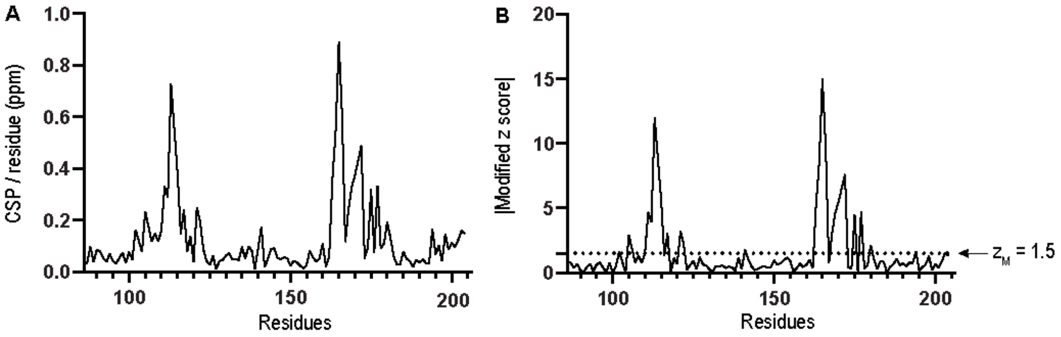


**Figure S3.** NMR chemical shift perturbation (CSP) between 1918 and PR8 NS1s. (**A**) NMR CSP values vs. residue. The CSP values were calculated by averaging the chemical shifts of all assigned 1H and 13C atoms in backbone and side chains. (**B**) Modified z-score ($z_M$) caclulated using NMR CSP data. A significant CSP is defined as a its $z_M$ greater than 1.5 (shown in dotted line).

Figure S4

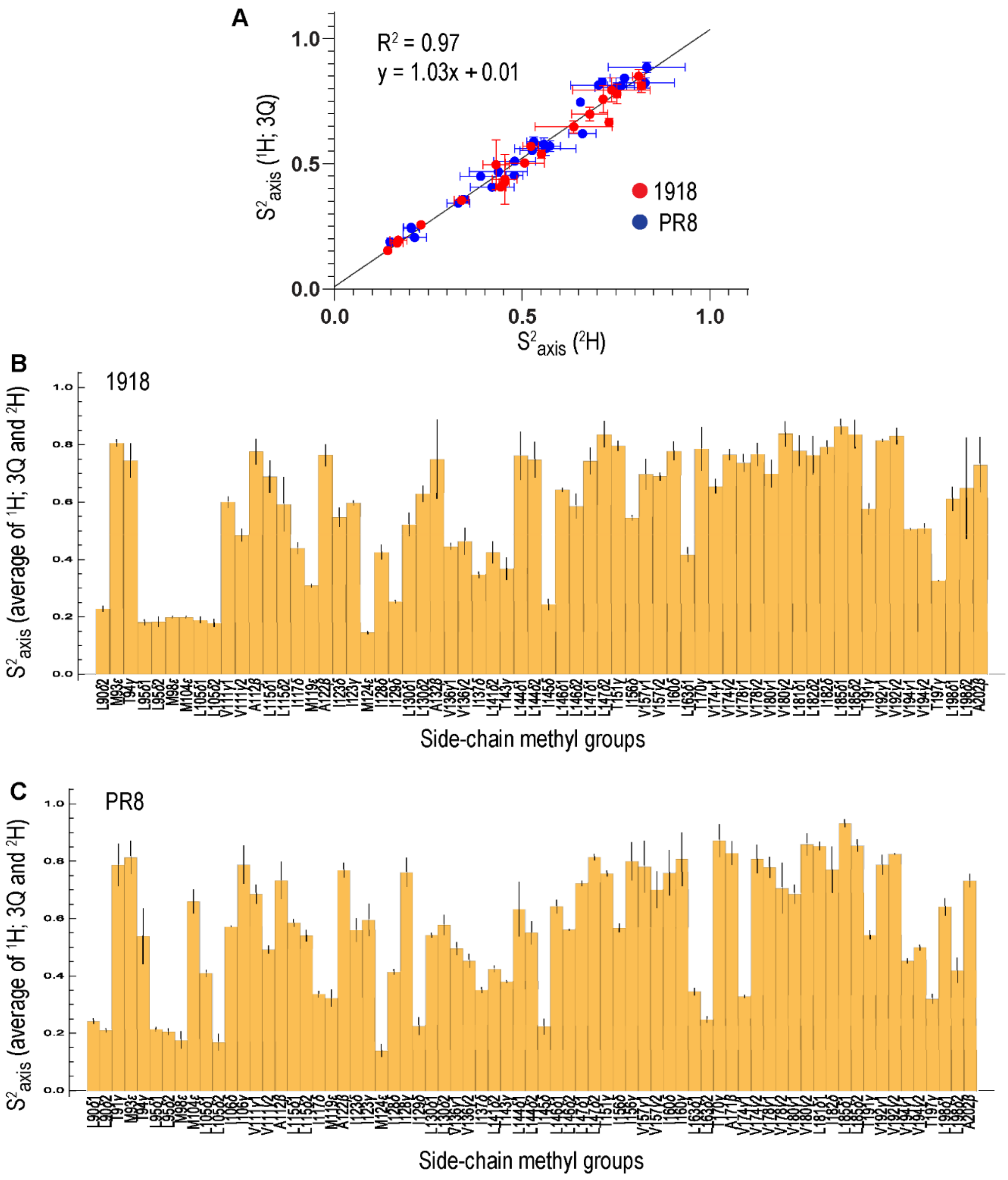


**Figure S4.** Comparison of NMR $S^2_{axis}$ values. (**A**) Correlations between $S^2_{axis}$ values of I, L, and V residues measured by $^2H$ relaxation and intramethyl $^1H$-$^1H$ cross-correlated relaxation (CCR) using forbidden triple quantum (3Q) approaches. The black line represents the global linear regression of 1918 and PR8 NS1 data. Plot of averaged $S^2_{axis}$ vs. methyl-bearing residues (A, I, L, M, V, and T) for (**B**) 1918 and (**C**) PR8 NS1s. $S^2_{axis}$ values of I, L, and V residues were averaged when both $^2H$-derived and CCR-derived values are avaialable.

Figure S5

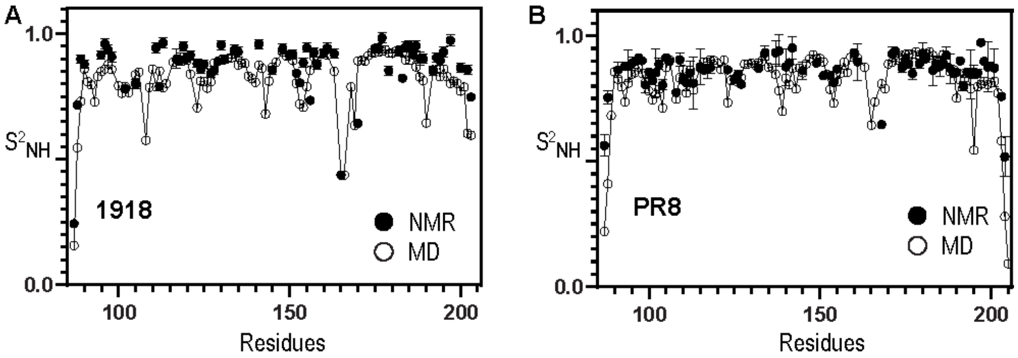


**Figure S5.** Backbone order parameters ($S^2_{NH}$). Comparison of $S^2_{NH}$ values for (**A**) 1918 and (**B**) PR8 NS1s, measured using NMR (closed circles) and MD (open circles connected by lines) approaches.

Figure S6

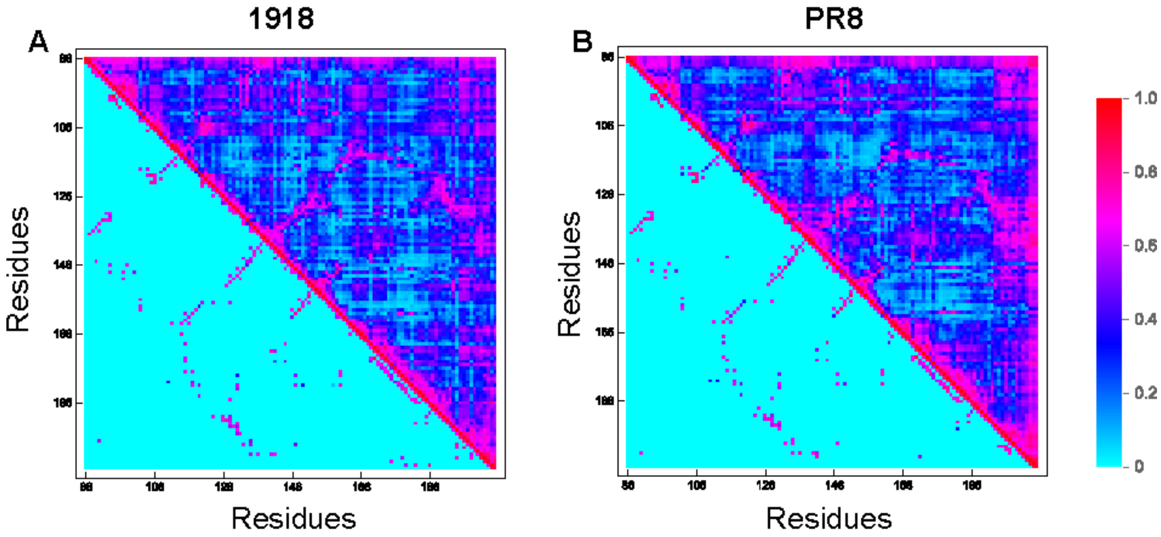


**Figure S6.** Correlated dynamics in NS1. Comparison of generalized cross-correlation (GCC) (upper triangle) and filtered GCC (lower triangle) for (**A**) 1918 and (**B**) PR8 NS1s. Network analysis was perfromed using the filtered GCC.

Figure S7

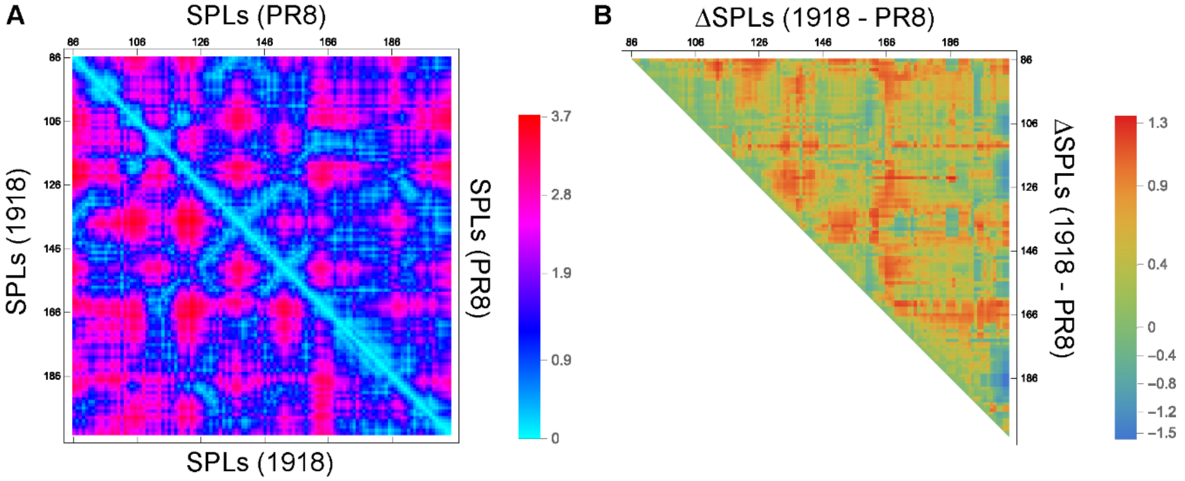


**C**

| | | **PR8** | | | |
|---|---|---|---|---|---|
| | | **103** | **106** | **114** | **171** |
| **1918** | **103** | | 0.85 | 2.12 | 2.21 |
| | **106** | 0.93 | | 2.24 | 2.31 |
| | **114** | 2.32 | 2.34 | | 0.60 |
| | **171** | 2.51 | 2.70 | 1.31 | |

**D**

| | **1918** | | | | |
|---|---|---|---|---|---|
| | **89** | **95** | **98** | **145** | **146** |
| **103** | 2.22 | 1.82 | 1.83 | 1.77 | 1.29 |
| **106** | 2.48 | 2.08 | 2.09 | 2.03 | 1.65 |
| **114** | 2.46 | 2.22 | 2.17 | 2.01 | 1.73 |
| **171** | 1.54 | 2.33 | 2.17 | 1.51 | 1.73 |

| | **PR8** | | | | |
|---|---|---|---|---|---|
| | **89** | **95** | **98** | **145** | **146** |
| **103** | 2.10 | 1.71 | 1.76 | 1.72 | 1.36 |
| **106** | 2.38 | 1.99 | 2.04 | 2.00 | 1.64 |
| **114** | 1.51 | 1.91 | 1.73 | 1.13 | 1.33 |
| **171** | 1.45 | 1.87 | 1.69 | 1.07 | 1.30 |

**Figure S7.** Network analysis of NS1s. (**A**) Comparison of the shortest path length (SPL) between nodes in 1918 NS1 (lower triangle) and PR8 NS1 (upper triangle). (**B**) The differences in SPL (ΔSPLs) between 1918 and PR8 NS1s. (**C**) SPLs between mutation sites in 1918 (lower triangle) and PR8 (upper triangle) NS1s. (**D**) SPLs between mutation sites and the core p85β-binding residues.

Figure S8

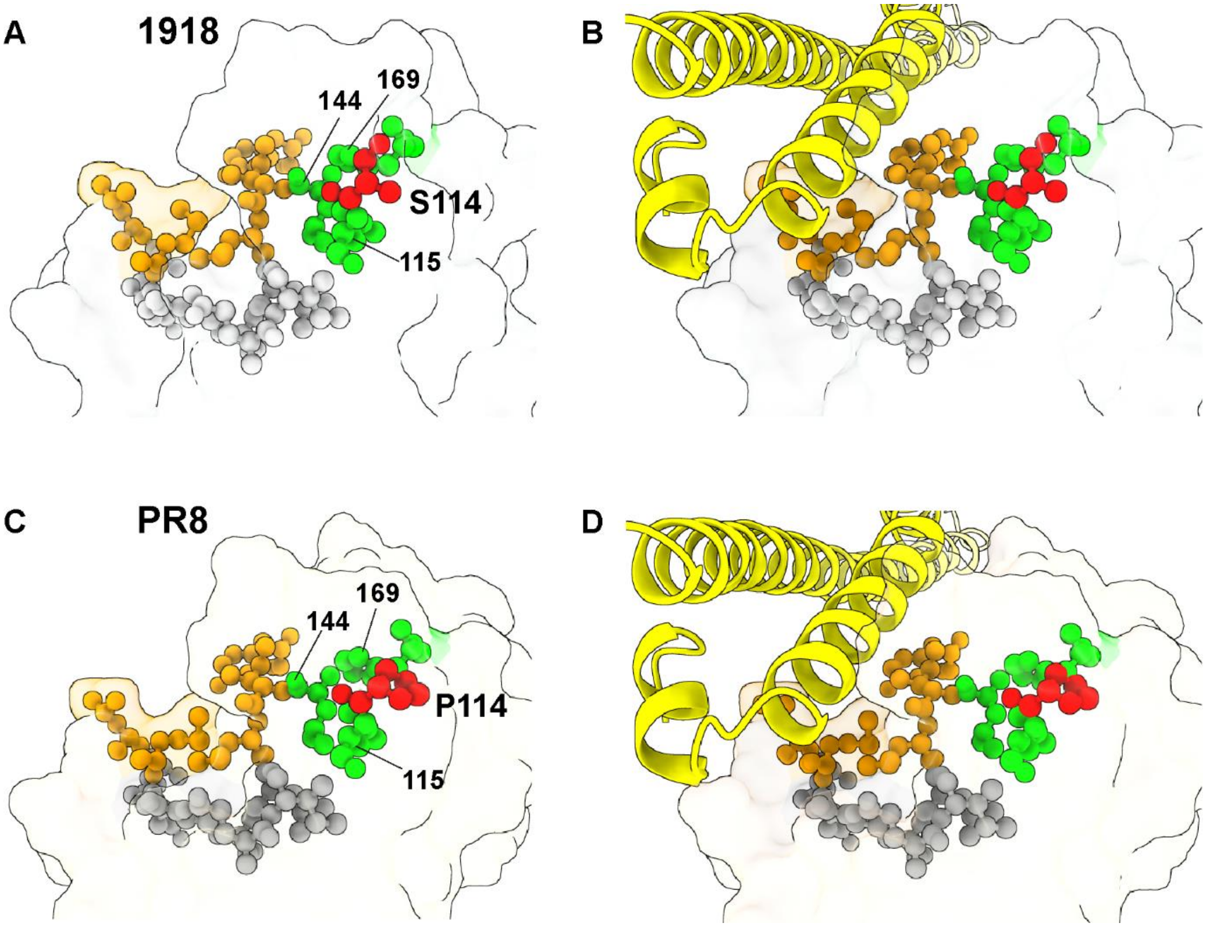


**Figure S8.** The shortest path (SP) between residues 114 (shown in red) and core p85β-binding residues 89, 95, 98, 145, and 146 (shown in orange). All residues involved in the SPs are depicted as a ball-and-stick model. SPs between residue 114 and core p85β-binding residues in panels (**A**) and (**C**) for NS1s and (**B**) and (**D**) for NS1s with bound p85β (depicted in yellow cartoon representation). Residues with significant NMR CSP or $\Delta S^2_{axis}$ are highlighted in lime with their residue numbers. Other residues in the pathways are shown as gray ball-and-stick model.

Figure S8 (continued)

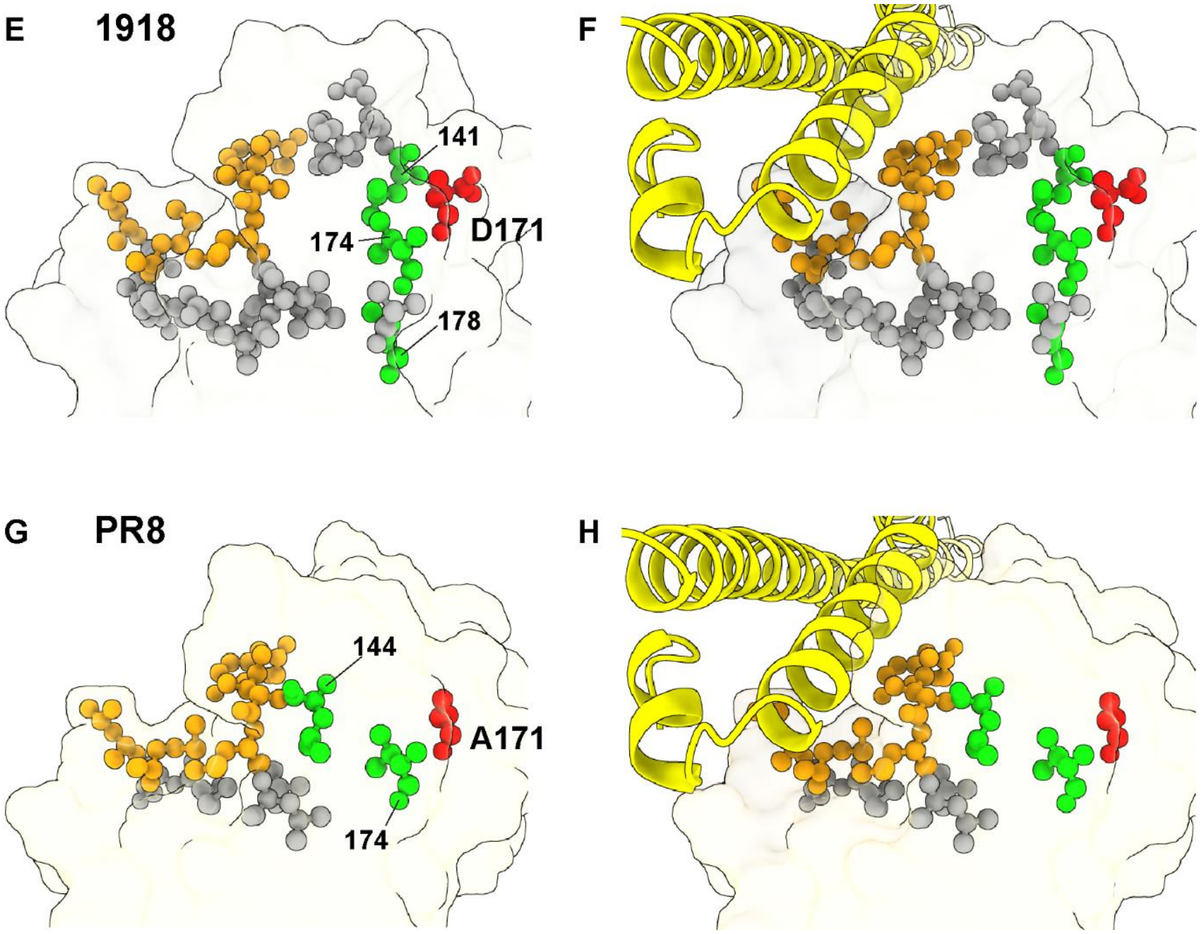


**Figure S8 (continued).** The shortest path (SP) between residues 171 (shown in red) and core p85β-binding residues 89, 95, 98, 145, and 146 (shown in orange). All residues involved in the SPs are depicted as a ball-and-stick model. SPs between residue 114 and core p85β-binding residues in panels (**A**) and (**C**) for NS1s and (**B**) and (**D**) for NS1s with bound p85β (depicted in yellow cartoon representation). Residues with significant NMR CSP or $\Delta S^2_{axis}$ are highlighted in lime with their residue numbers. Other residues in the pathways are shown as gray ball-and-stick model.

Table S1

**Table S1.** Communities identified in 1918 and PR8 NS1s.

| Communities | 1918 NS1 | PR8 NS1 |
|---|---|---|
| 1 (Cyan) | 110, 111, 112, 113, 114, 115, 116, 117, 141, 144, 160, 161, 162, 163, 164, 165, 166, 167, 168, 169, 170, 171, 172, 173, 174, 175, 176, 177, 180 | 111, 113, 114, 115, 116, 160, 162, 163, 164, 165, 166, 167, 168, 169, 170, 172, 173, 174, 175, 176, 177, 179 |
| 2 (Orchid) | 86, 87, 88, 89, 90, 91, 92, 93, 131, 132, 133, 134, 135, 136, 137, 138, 139, 140, 142, 143, 145, 146, 196, 197, 198, 199, 200, 201 | 86, 87, 88, 89, 90, 135, 136, 137, 138, 139, 140, 141, 142, 143, 171, 201, 202, 203, 204, 205 |
| 3 (Blue) | 126, 127, 128, 129, 130, 147, 178, 179, 182, 183, 185, 186, 187, 188, 189, 190, 191, 192, 193, 194, 195, 202, 203, 204, 205 | 123, 124, 125, 126, 127, 128, 129, 130, 147, 149, 178, 181, 182, 183, 185, 186, 187, 188, 189, 190, 191, 192, 193, 194 |
| 4 (Purple) | 102, 103, 104, 105, 106, 107, 108, 109, 118, 119, 120, 121, 122, 123, 124, 125, 149, 156, 157, 158, 159, 181, 184 | 102, 103, 104, 105, 106, 107, 108, 109, 110, 112, 117, 118, 119, 120, 121, 122, 156, 157, 158, 159, 180, 184 |
| 5 (Goldenrod) | 94, 95, 96, 97, 98, 99, 100, 101, 148, 150, 151, 152, 153, 154, 155 | 91, 92, 93, 94, 95, 96, 97, 98, 99, 100, 101, 131, 132, 133, 134, 144, 145, 146, 148, 150, 161, 197, 198 |
| 6 (Salmon) | | 151, 152, 153, 154, 155 |
| 7 (Dark Orchid) | | 195, 196, 199, 200 |

The name of colors in parentheses correspond to the colors used in Figure 5.
Communties 6 and 7 in PR8 NS1 are closely located with communities 2 and 5, respectively.
